# Andrographolide prevents necroptosis by suppressing generation of reactive oxygen species

**DOI:** 10.1101/2025.04.07.647496

**Authors:** Na Lu, Qing Li, Ling-han Duan, Rong Xu, Ya-ping Li, Fu-li Shi, Zhi-ya Zhou, Ying-qing Gan, Bo Hu, Jin-hua Li, Xian-hui He, Dong-yun Ouyang, Qing-bing Zha

## Abstract

Andrographolide (Andro), a natural product extracted from the Chinese traditional medicine herb *Andrographis paniculata*, has been applied for the treatment of diverse inflammatory diseases. However, its effects on necroptosis, a lytic form of cell death implicated in various inflammatory diseases, remain uncharacterized. In the current research, we investigate whether Andro and its derivatives can suppress necroptosis. The results demonstrate that Andro notably inhibits the necroptosis i<u>n</u> *in vitro* cellular models induced by either lipopolysaccharide (LPS) plus IDN-6556 or a combination of TNF-α, LCL-161 (Smac mimetic) and IDN-6556. In these cellular models, Andro exhibits inhibitory effects on the phosphorylation of receptor-interacting protein kinase 1 (RIPK1), RIPK3, and mixed lineage kinase domain-like protein (MLKL), as well as on the formation of necrosomes. Specifically, Andro reduces intracellular reactive oxygen species (ROS) and mitochondrial superoxide (mtROS), preserves mitochondrial membrane potential during necroptotic induction, and activates the antioxidant transcription factor nuclear factor E2-related factor 2 (Nrf2). Upon the necroptotic stimulation, some mitochondrial proteins such as Bcl-2 and Bak oligomerize and colocalize with RIPK1, RIPK3, and phosphorylated MLKL (p-MLKL) in necrosomes. However, such a process of necrosome formation can be prevented by Andro. In contrast, derivatives including dehydroandrographolide, neoandrographolide, 14-deoxy-11,12-didehydroandrographolide, and 14-deoxyandrographolide show no anti-necroptotic effects and fail to upregulate Nrf2. Collectively, our findings demonstrate that Andro specifically inhibits the RIPK1/RIPK3/MLKL signaling axis to suppress necroptosis, highlighting its therapeutic potential against necroptosis-related disorders.

## 1. Introduction

*Andrographis paniculata,* a Chinese traditional medicine, is widely used to treat inflammatory diseases including gastroenteritis, tracheitis, lung abscess, cholecystitis, and oropharyngeal swelling/pain. Andrographolide (Andro), a diterpene lactone compound isolated from this plant, has demonstrated therapeutic efficacy against a number of inflammatory conditions such as pneumonia ^[1]^, colitis ^[2, 3]^, myocarditis ^[4]^, and rheumatoid arthritis ^[5]^.

Previous studies have revealed multiple mechanisms underlying the anti-inflammatory effects of Andro. First, Andro covalently modifies the cysteine residues within the nuclear factor (NF)-κB p50 subunit to suppress NF-κB signaling ^[6, 7]^. Second, Andro can activate the nuclear factor E2-related factor 2 (Nrf2) ^[8]^. Third, Andro can inhibit inflammasome activation, thus preventing pyroptosis ^[9, 10]^. Despite these advances, more investigations are warranted for comprehensive understanding of its anti-inflammatory mechanisms.

Regulated cell death (RCD) is a key event in inflammatory organ injury. Among various forms of RCD, necroptosis exerts a critical influence on modulating physiological and pathological processes, causing a range of human diseases, including ischemic brain injury, immune system diseases, and cancer ^[11]^. This lytic cell death is characterized by the disruption of the cell membrane, resulting in the release of cellular components and inflammatory mediators, consequently triggering inflammation. Unlike other forms of RCD, necroptosis is independent of cysteinyl aspartate-specific proteinases (caspases) but is controlled by the receptor-interacting protein kinase 1 (RIPK1)/RIPK3/mixed lineage kinase domain-like protein (MLKL) signaling pathway. Mechanistically, phosphorylated RIPK1 recruits and phosphorylates RIPK3 ^[12, 13]^, thus forming a necrosome complex that subsequently phosphorylates/activates MLKL. Activated MLKL translocates to the plasma membrane and oligomerizes there to form membrane-disrupting pores ^[14]^, which can be inhibited by by Necrostatin-1 (Nec-1), a specific RIPK1 inhibitor ^[15]^. Such a necroptotic signaling pathway can be initiated by the binding of tumor necrosis factor (TNF) to its receptor TNFR1, and viral nucleic acids, *etc.* ^[16, 17]^. Therefore, targeting necroptosis appears an effective and feasible approach for treating such diseases.

In this study, we intended to investigate Andro’s potential to inhibit necroptosis as a novel anti-inflammatory mechanism. Our evidence shows that Andro effectively attenuated the necroptosis induced by TNF-α/LCL-161 (Smac mimetic)/IDN-6556 (a pan-caspase inhibitor) or LPS/IDN-6556. Mechanistically, Andro inhibited the activation of the RIPK1-RIPK3-MLKL signaling pathway, while it decreased intracellular reactive oxygen species (ROS) accumulation and mitochondrial superoxide (mtROS) production, thereby preserving mitochondrial membrane potential (MMP). Notably, four structural derivatives of Andro lacked comparable inhibitory activities. *In vivo*, Andro could alleviate the uterine injury induced by TNF-α in C57BL/6J mice. These findings propose Andro as a promising therapeutic candidate for necroptosis-mediated inflammatory diseases.

## 2. Materials and methods

### Reagents and antibodies

Andrographolide (A823169) was purchased from Macklin (Shanghai, China) and dissolved in dimethyl sulfoxide (DMSO). Andrographolide (B20207), 14-deoxy-11,12-didehydroandrographolide (B30518), 14-deoxyandrographolide (B24387), neoandrographolide (B21392), and dehydroandrographolide (B21275) were purchased from Yuanye (Shanghai, China). Lipopolysaccharide (LPS, Escherichia coli O111:B4), Hoechst 33342, propidium iodide (PI), DMSO, and DL-dithiothreitol (DTT) were all purchased from Sigma-Aldrich (St Louis, USA). IDN-6556 (S7775) and LCL-161 (S7009) were purchased from Selleck Chemicals (Houston, TX, USA). Fetal bovine serum (FBS), Dulbecco’s Modified Eagle’s Medium (DMEM) with high glucose, streptomycin, and penicillin were purchased from ThermoFisher (Carlsbad, CA, USA). Specific antibodies against β-actin (#3700), phospho(p)-RIPK1 (#53286), p-RIPK3 (#91702), p-MLKL (#37333), RIPK1 (#3493), RIPK3 (#95702), MLKL (#37705), horseradish peroxidase (HRP)-conjugated horse-anti-mouse IgG (#7076), and HRP-conjugated goat-anti-rabbit IgG (#7074) were purchased from Cell Signaling Technology (Danvers, MA, USA).

### Cell Culture

The murine macrophage cell line J774A.1 was obtained from the Kunming Cell Bank, CAS (Kunming, China) and cultured in complete DMEM supplemented with 10% FBS, 100 μg/ml streptomycin, and 100 U/ml penicillin). For the experiments, J774A.1 cells were seeded at a density of 8 × 10^4^ cells/well in a 24-well plate or 3.2 × 10^5^ cells/well in a 6-well plate. Human HT-29 colorectal adenocarcinoma cells were obtained from the Cell Bank of the Chinese Academy of Sciences (Shanghai, China) and cultured in complete McCoy’s 5A medium. For the experiments, HT-29 cells were seeded at a density of 1.4 × 10^5^ cells/well in a 24-well plate.

Bone marrow-derived macrophages (BMDMs) were obtained as previously reported ^[18]^. Briefly, the bone marrow cells were harvested from the hind femora and tibias of mice and seeded in a 10-cm petri dish containing 10 ml of BM-Mac medium (containing 80% complete DMEM and 20% M-CSF-conditioned medium derived from L-929 cells). After 3 days, supplemented each dish with 5 ml of BM-Mac medium and continued to culture the cells for another 3 days. For the experiments, BMDMs were seeded in a 24-well plate (2.5 × 10^5^ cells) or a 6-well plate (1.6 × 10^6^ cells). All cells were incubated in a humidified incubator with 5% CO_2_ at 37 °C.

### Induction of necroptosis

Necroptosis was induced, as reported previously ^[19]^. A combination of TNF-α (T, 30 ng/ml), LCL-161 (S, 10 μmol/l for BMDMs and J774A.1 cells or 50 μmol/l for HT-29 cells) and IDN-6556 (I, 30 μmol/l) (hereafter referred to as TSI), or a combination of LPS (L, 0.5 μg/ml) and IDN-6556 (I, 30 μmol/l) (hereafter referred to as LI) was used for indicated time length.

### Cell death assay

Cell death was evaluated by a PI incorporation assay as previously described ^[20]^. After the indicated treatments, the cells were stained with a solution containing 2 μg/ml PI and 5 μg/ml Hoechst 33342 in sterile PBS in the dark for 10 min. Then, the cells were observed under a Zeiss Axio Observer D1 microscope (Carl Zeiss, Gottingen, Germany), and the images were captured.

### Western blot analysis

Western blotting was performed as previously described ^[21]^. Proteins in cell lysates were separated by sodium dodecyl sulfate-polyacrylamide gel (SDS-PAGE) electrophoresis and subsequently transferred to PVDF membranes (03010040001; Roche, Mannheim, Germany). The membranes were then blocked with a blocking buffer for 1 h. After that, the membranes were incubated with indicated antibody at 4° C overnight. Subsequently, the membranes were incubated with a suitable HRP-conjugated antibody and then treated with an enhanced chemiluminescence kit (BeyoECL Plus; Beyotime, Shanghai, China) to visualize the target bands, which were then captured on X-ray films (Carestream, Xiamen, China).

### Immunofluorescence microscopy

Immunofluorescence (IF) analysis was performed as previously described ^[22]^. Briefly, BMDMs were seeded in glass-bottom dishes (2.5×10^5^ cells). After the indicated treatments, the cells were fixed with 4% paraformaldehyde in PBS for 15 min, followed by being permeabilized in cold methanol at −20 [for 10 min. Next, the cells were then blocked with a blocking solution (5% goat serum in PBS) for 1 h and then incubated with indicated antibodies (1:300) at 4 °C overnight. After that, the cells were stained with CF568-conjugated goat-anti-rabbit IgG (#SAB4600084, Sigma-Aldrich) (1:300) and CF488-conjugated goat-anti-mouse IgG (#SAB4600237, Sigma-Aldrich) (1:300) for 1 h. Subsequently, the cells were stained in PBS solution containing Hoechst 33342 for 10 min. Finally, Cell images were captured using the Axio Observer D1 microscope.

### Detection of intracellular ROS and mitochondrial superoxide

As previously described ^[23]^, an ROS assay kit (S0033; Beyotime) was utilized to evaluate intracellular ROS levels, while MitoSOX red (M36008, ThermoFisher) was used to detect the levels of mitochondrial superoxide (mtROS). After the indicated treatments, cells were assayed using the above reagents in an incubator for appropriate time lengths following the instructions of the suppliers, respectively. Finally, the fluorescence of DCFH-DA or MitoSOX within the cells was examined using the Axio Observer D1 microscope.

### Detection of MMP

MMP was determined using a TMRE assay kit (C2001, Beyotime) and JC-1 probe (C2006, Beyotime), respectively. After the indicated treatments, cells were incubated with 1 µg/ml TMRE or 5 µg/ml JC-1 staining solution following the instructions of the suppliers. The intracellular red fluorescence of TMRE was observed using the Axio Observer D1 microscope, and the JC-1 staining was analyzed using a flow cytometer (Attune NxT, Thermo Fisher).

### TNF-**α**-induced systemic inflammatory response syndrome (SIRS) model

Female C57BL/6 mice were randomly assigned to 4 groups (n = 5 per group): Vehicle group, Andro group, TNF-α group, and Andro + TNF-α group. The mice in TNF-α and Andro + TNF-α groups were intravenously (*i.v.*) injected with a single dose of TNF-α (5 μg per mouse), while vehicle group was intravenously injected with sterile PBS. The mice in Andro and Andro + TNF-α groups were intraperitoneally (*i.g.*) administered with Andro (0.5 mg/kg or 1 mg/kg body weight) thrice (at 1 h before, and 2 h and 5 h after TNF-α injection, respectively). Nine hours after TNF-α administration, the mice were anesthetized and sacrificed by cervical dislocation, followed by collection of uterus tissues.

### Statistical analysis

Data were presented as the mean ± standard deviation (SD) and were analyzed for statistical significance using GraphPad Prism 7.0 software (GraphPad Software Inc, San Diego, CA, USA). The statistical significance among multiple groups was determined using a one-way analysis of variance (ANOVA) followed by a Tukey post hoc test. Statistical significance was defined as: \**P* < 0.05, \*\**P* < 0.01, and \*\*\**P* < 0.001.

## 3. Results

### 3.1 Andro inhibits TSI- or LI-induced necroptosis in macrophages and HT-29 cells

The anti-necroptotic effect of Andro was evaluated in macrophages and HT-29 cells treated with TSI or LI. Both TSI- and LI-treated cells exhibited characteristic necroptotic features, including cellular swelling and membrane rupture ^[24]^. As indicated by PI incorporation, Andro dose-dependently inhibited the necroptosis in J774A.1 macrophages **(Fig. 1A-D)**, HT-29 adenocarcinoma cells **(Fig. 1E, F)**, and mouse BMDMs (**Supplementary Fig. S1A-D**). These findings demonstrate that Andro effectively inhibits necroptosis in both macrophages and other cell types.

**Fig. 1.**
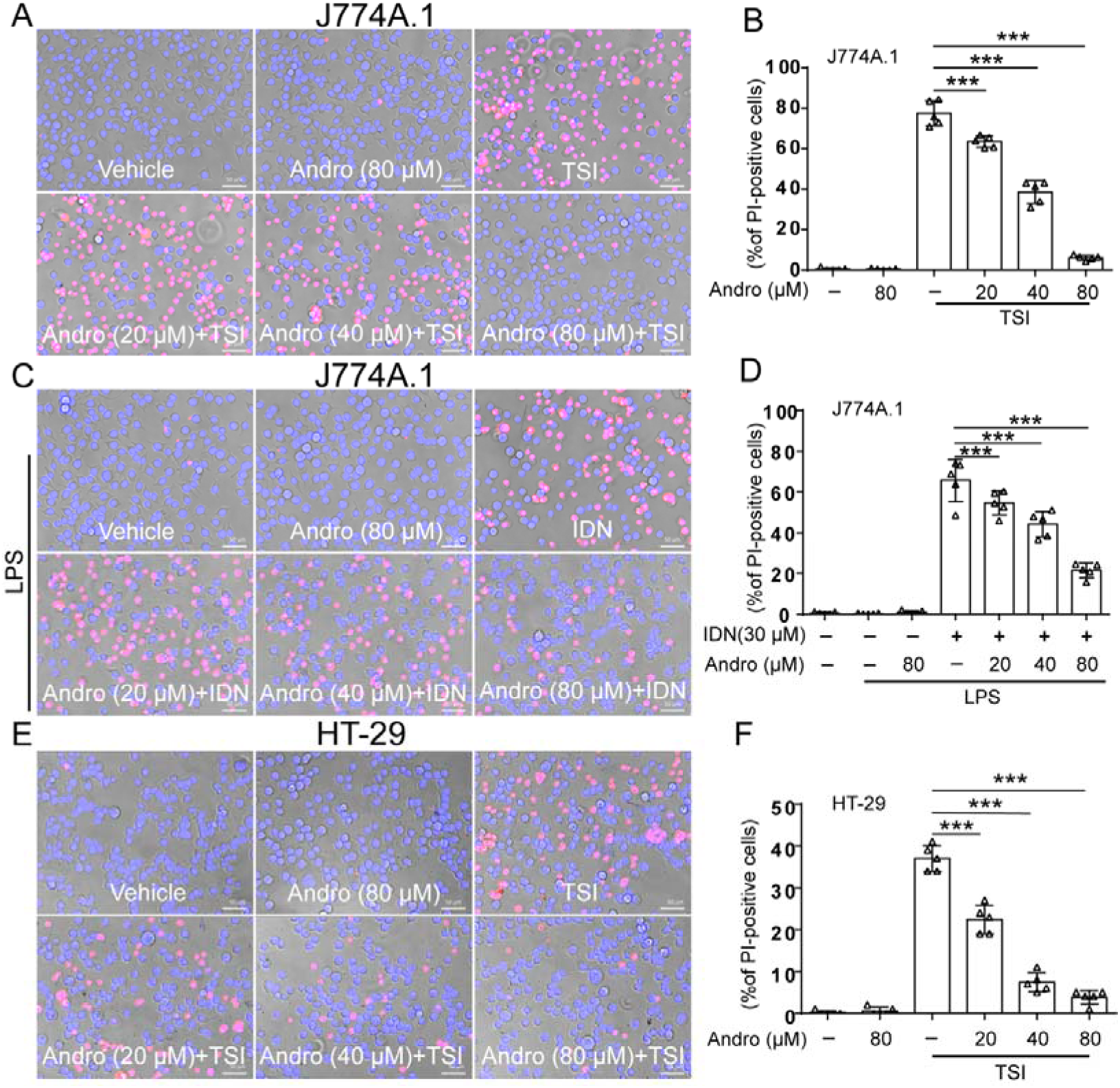
Andro suppresses necroptosis in macrophages and HT-29 cells. J774A.1 cells and HT-29 cells were pretreated with or without Andro for 1 h, followed by stimulation with the combination of TNF-α, LCL-161, and IDN-6556 (TSI) (A, B, E, F), or LPS plus IDN-6556 (LI) (C, D) in the presence or absence of Andro for 2 h. The ratios of cell death were evaluated through staining cells with PI (red, cells undergoing lytic cell death) and Hoechst 33342 (blue, all nuclei). Scale bars, 50 μm. Data are shown as mean ± SD (n = 5). *\*P* < 0.05; *\*\*\*P* < 0.001; ns, not significant.

### 3.2 Andro suppresses the necroptotic signaling pathway in mouse macrophages

Next, we investigated whether Andro inhibited the necroptotic signaling pathway activation in TSI- or LI-stimulated macrophages. Western blot analysis showed that after TSI or LI treatment, the levels of phosphorylated RIPK1, RIPK3, and MLKL were profoundly increased in BMDMs (**Fig. 2A-H**) and J774A.1 cells (**Supplementary Fig. S2A-H**), confirming activation of the necroptotic signaling pathway. Andro treatment significantly attenuated these phosphorylation events (**Fig. 2; Supplementary Fig. S2**), paralleling its inhibition of cell death induced by TSI or LI. These findings indicate that Andro blocks necroptosis by inhibiting RIPK1/RIPK3/MLKL signaling.

**Fig. 2.**
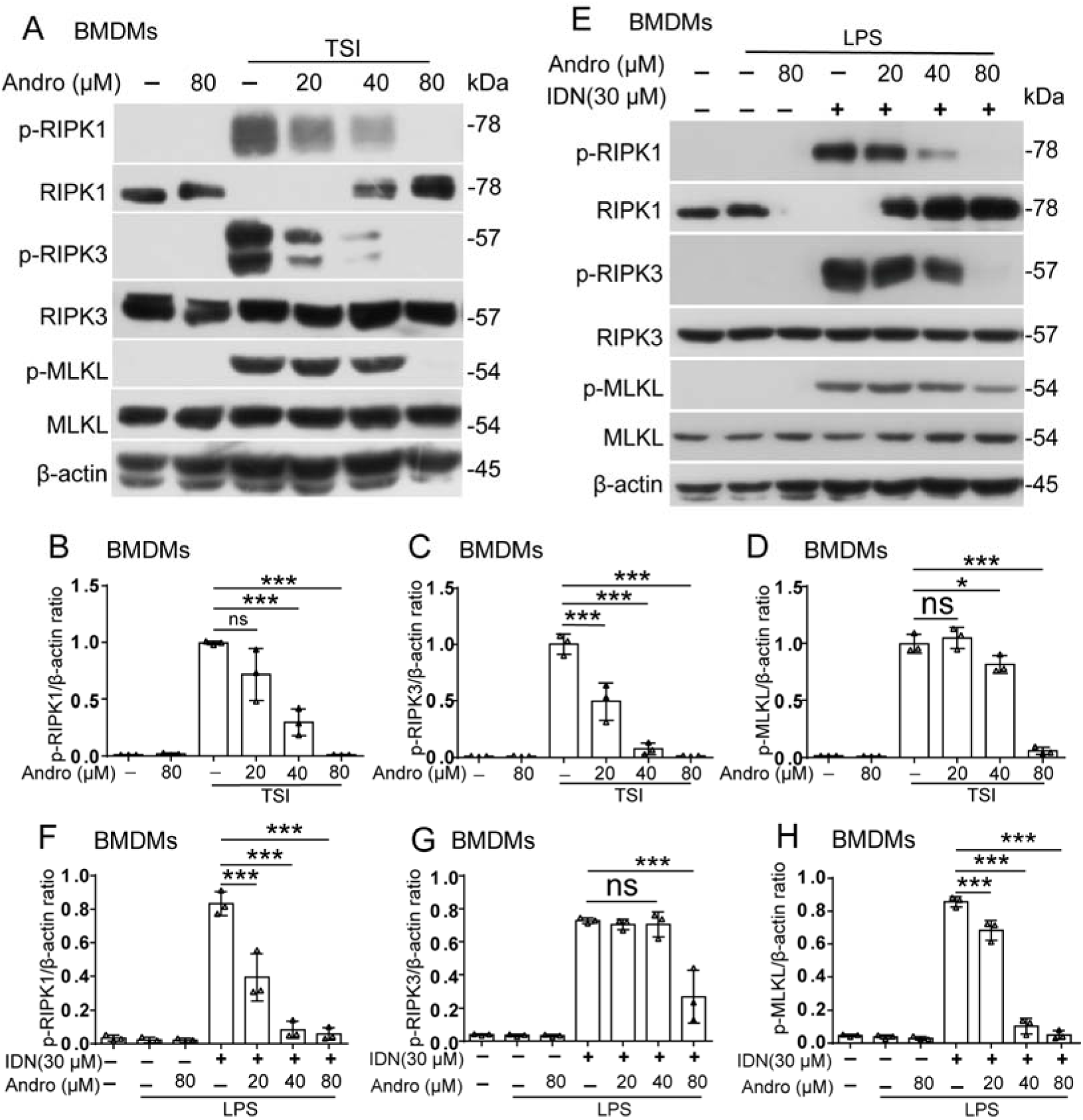
Andrographolide suppresses the necroptotic signaling pathway. BMDMs were treated with TNF-α, LCL-161, and IDN-6556 (TSI) (**A-D**), or LPS plus IDN-6556 (LI) (**E-H**) as described in Fig. 1. Western blotting was used to detect the proteins of the necroptotic pathway. The loading control was β-actin. Relative levels of indicated proteins to that of β-actin in cell lysates were determined, with the levels of TSI or LI group being set as 1.0, respectively. Data are shown as mean ± SD (n = 3). *\*P* < 0.05; *\*\*\*P* < 0.001; ns, not significant.

### 3.3 Andro blocks the formation of RIPK1/RIPK3 necrosome induced by TSI or LI

In normal cells, RIPK1 and RIPK3 are diffusely and evenly distributed in the cytoplasm. However, upon necroptotic stimulation, RIPK1 and RIPK3 aggregate into punctate necrosomes in the cytoplasm, with subsequent recruitment and activation of MLKL ^[18]^. In this study, IF analysis was used to investigate whether Andro could inhibit the necrosome formation in both TSI-**(Fig. 3A)** and LI-**(Fig. 3B)** treated BMDMs. It was observed that RIPK1 (red) or RIPK3 (green) formed aggregates in the cytoplasm, but Andro pretreatment abolished their co-localization. However, such co-localization of RIPK1 and RIPK3 aggregates was not observed in the control, Andro, and Andro+TSI/LI groups. This disruption of necrosome formation by Andro mechanistically explains its anti-necroptotic activity.

**Fig. 3.**
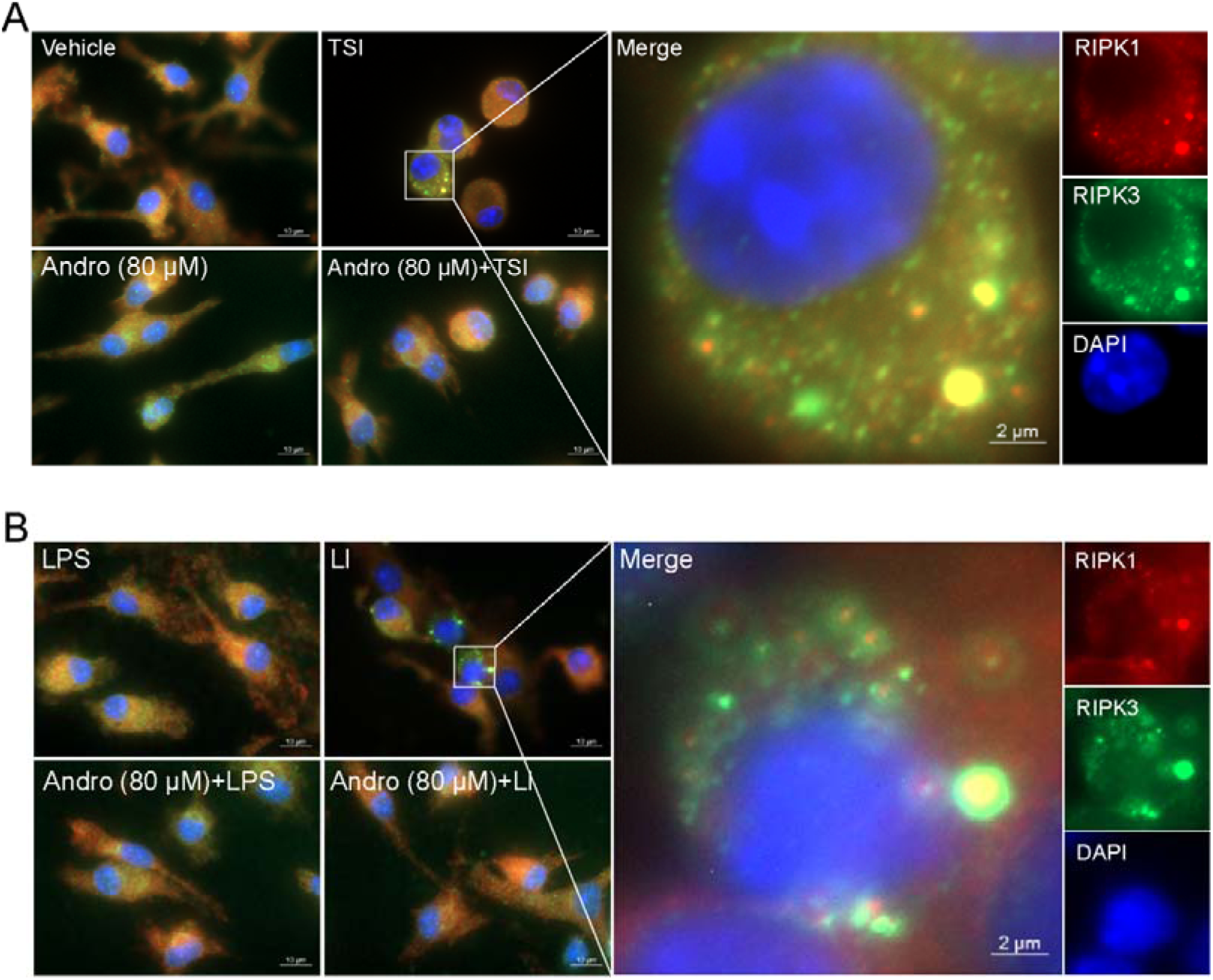
Andrographolide prevents the formation of necrosome in macrophages stimulated with TSI or LI. BMDMs were treated as described in Fig. 1. The immunofluorescence technique was used to observe the distribution and co-localization of RIPK1 (red) and RIPK3 (green). Nuclei were stained with Hoechst 33342 (blue). Scale bar, 10 µm; inset scale bar, 2 µm.

### 3.4 Andro scavenges TSI-induced ROS and prevents ROS-mediated mitochondrial protein oligomerization, thus reducing mitochondrial damage

Reciprocal regulation between ROS and necroptotic signaling has been suggested by emerging evidence: (1) RIPK1 senses ROS via its three key cysteine residues, leading to its autophosphorylation at Serine 161, and subsequently recruiting RIPK3 to form a functional necrosome ^[25]^; (2) ROS mediates the cross-linking of MLKL proteins through disulfide bonds, facilitating their oligomerization and necroptosis execution ^[26]^; (3) mtROS not only triggers pyroptosis-mediated mtDNA release but also activates the necroptotic signaling pathway ^[27]^; (4) activated RIPK1 stimulates the generation of ROS, establishing a pathogenic feedforward loop ^[28]^. Therefore, we next sought to investigate whether Andro could suppress the generation of intracellular ROS and mtROS in TSI-stimulated cells. DCFH-DA was used as a probe for intracellular ROS, and MitoSOX was used to detect the mitochondrial superoxide levels in TSI-activated cells. The results showed that Andro significantly suppressed both intracellular ROS (**Fig. 4A**, **B**) and mtROS **(Supplementary Fig. S3A, B)**, suggesting that Andro inhibits necroptosis by inhibiting ROS production.

**Fig. 4.**
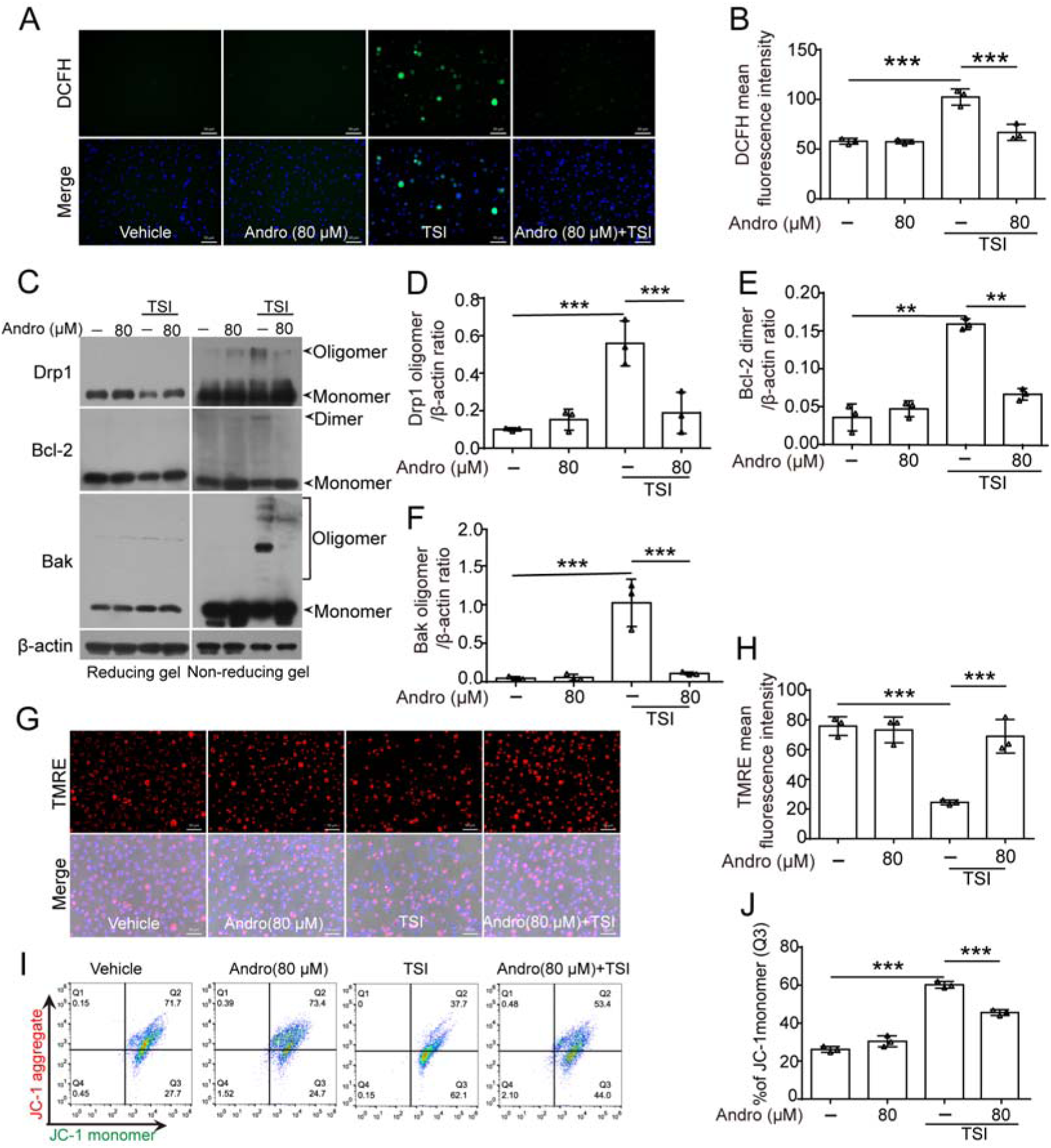
Andrographolide suppresses the production of ROS induced by TSI-induced in BMDMs. BMDMs were treated as described in Fig. 1. (**A**) After staining with DCFH-DA, ROS (green) was observed using a fluorescence microscope. (**B**) Quantitative analysis of DCFH fluorescence intensity in (**A**). (**C**) Western blotting analysis was performed to examine the expression of indicated proteins after reduced and non-reduced electrophoresis. (**D-F**) Relative levels of oligomerized Drp1, Bcl-2, and Bak to that of β-actin in cell lysates were determined, with the levels of TSI group being set as 1.0, respectively. (**G**) After being stained with TMRE, the cells were observed by a fluorescence microscope. Nuclei were stained with Hoechst 33342 (blue). Scale bar, 50 μm. (**H**) Mitochondrial membrane potential (MMP) was evaluated as the ratio of TMRE-fluorescence-positive cells. (**I**) MMP was analyzed by flow cytometry after the cells were stained with JC-1 reagent. (**J**) Loss of MMP was calculated as the ratios of cells with JC-1 monomers in (**I**). Data are shown as mean ± SD (*n* = 3). *\*\*\*P* < 0.001.

Building on our prior findings that both glycometabolic and ROS stresses induce disulfide-linked oligomerization of certain mitochondrial proteins (Drp1, Bcl-2, and Bak) ^[21]^, we examined whether this would take place under necroptotic conditions. By non-reducing electrophoresis and Western blotting, oligomerized Drp1, Bcl-2, and Bak were observed in TSI-treated macrophages, which was abolished by Andro treatment (**Fig. 4C**-**F**), indicating ROS-dependent crosslinking of these proteins during necroptosis.

The ROS and oligomerized mitochondrial proteins induced by TSI are likely to cause mitochondrial damage. To substantiate this concept, the reduction of MMP induced by TSI treatment was detected using both the TMRE assay (**Fig. 4G**, **H**) and the JC-1 assay combined with flow cytometry analysis (**Fig. 4I**, **J**). TSI caused marked loss of MMP as evidenced by decreased TMRE fluorescence (**Fig. 4G**, **H**) and increased JC-1 green monomers (**Fig. 4I**, **J**), which was reversed by Andro. These data collectively indicate that Andro preserves mitochondrial function by preventing ROS accumulation and subsequent oxidative protein modifications.

### 3.5 Andro inhibits Bcl-2 and Bak being incorporated into the necrosome induced by TSI

Recent studies have indicated that RIPK1 and ROS-mediated transfer of MLKL, Bak, and Drp1 to mitochondria leads to mitochondrial damage and necroptosis ^[29]^. We supposed that the proteins of the necroptotic pathway might also be transferred to the mitochondria of TSI-treated macrophages in a similar manner to activate necroptosis. The distributions of these proteins were observed by IF microscopy. Our data showed that Bcl-2, Bak, and Tom20 (a mitochondrial marker) formed a speck in TSI-treated cells (**Fig. 5A-C**), which were colocalized with the RIPK1/RIPK3 aggregates (**Fig. 5D-E**), suggesting assembly of mitochondrial-associated necrosomes. Andro pretreatment abolished these interactions (**Fig. 5**). The formation of necrosomes comprising Bcl-2, Bak, RIPK1, and RIPK3 should have caused mitochondrial disruption as the expression of Tom20 was much lower in TSI group than in other groups (**Fig. 5B, C**). While our non-reducing gels demonstrated Bcl-2/Bak oligomerization (**Fig. 4C**), more investigations are warranted to determine whether these proteins in the necrosome <u>have</u> been cross-linked (oligomerized) through disulfide bonds. These findings indicate that ROS-induced mitochondrial protein aggregation may bridge the necroptotic signaling and organelle damage.

**Fig. 5.**
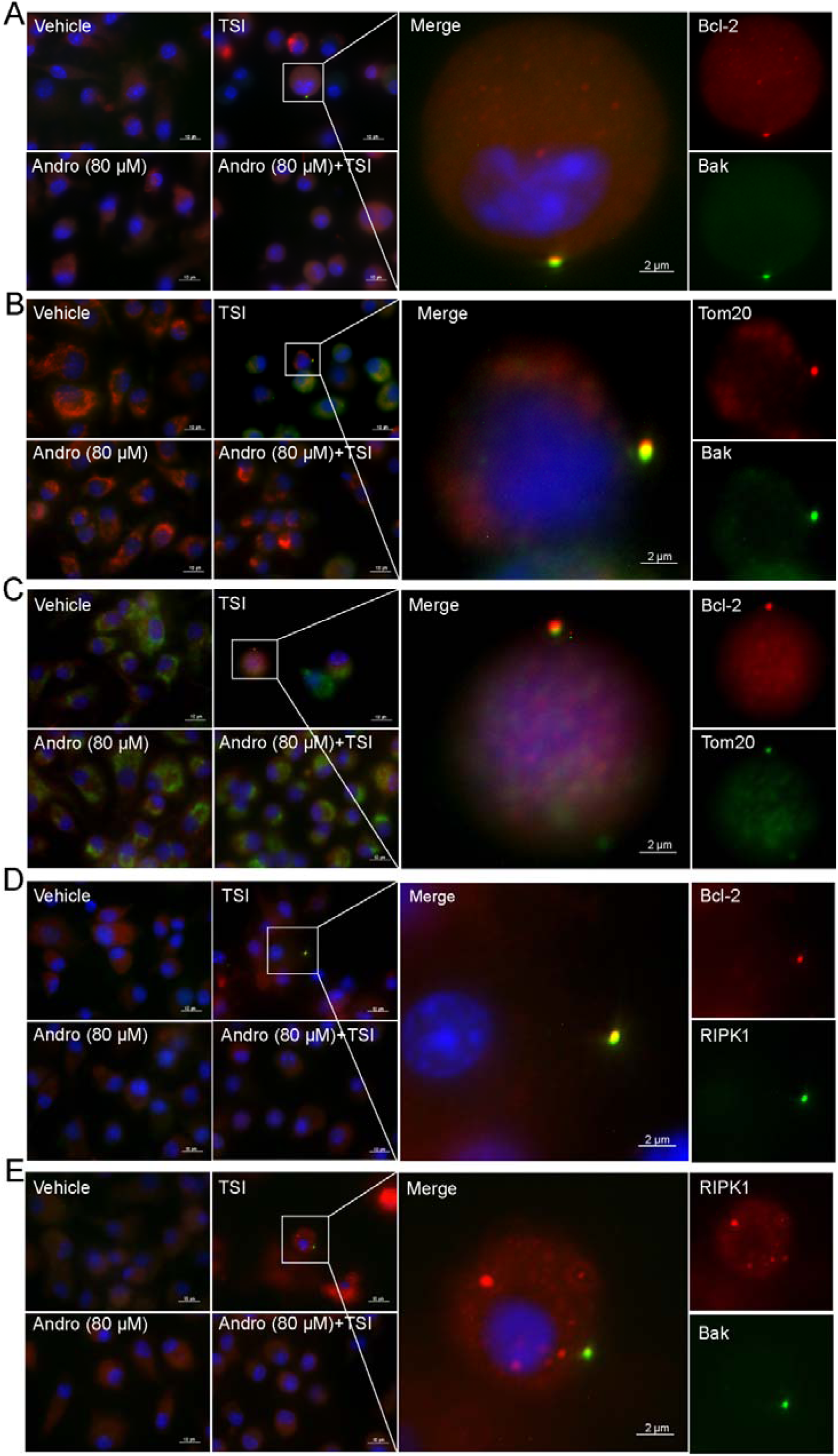
Andrographolide prevents the formation of necrosomes in TSI-induced BMDMs. BMDMs were treated with TSI as described in Fig. 1. The distribution of indicated proteins (green or red) was observed by immunofluorescence microscopy. Nuclei were stained with Hoechst 33342 (blue). Scale bar, 10 µm; inset scale bar, 2 µm.

### 3.6 Andro induces the activation of Nrf2-heme oxygenase-1 (HO-1) signaling pathway

HO-1, a cytoprotective enzyme regulated by Nrf2 through antioxidant response elements, plays a critical role in safeguarding cells against oxidative stress ^[30]^. Under normal conditions, Nrf2 maintains at a low level as it is sequestered in the cytoplasm by binding its cytoplasmic inhibitor KEAP1 and tends to be degraded through ubiquitin-proteasome pathway. Oxidative stress disrupts this interaction, enabling Nrf2 to translocate to the nucleus where it transcriptionally upregulates antioxidant-related proteins, including HO-1 ^[30]^. Building on prior evidence that Andro inhibits pyroptosis by promoting the expression of Nrf2 ^[31]^, we investigated whether Andro’s inhibitory effect on necroptosis depended on its regulation of Nrf2 expression. IF microscopy revealed that the translocation of Nrf2 to the nucleus was significantly increased after Andro treatment, irrespective of TSI treatment (**Fig. 6A)**. Consistent with this observation, western blotting analysis corroborated that Nrf2 levels were undetectable or minimal in the control and TSI-only groups, but were greatly increased in Andro-treated cells, regardless of TSI treatment (**Fig. 6B, C**). Consistent with Nrf2 activation, KEAP1 levels were reduced while HO-1 levels were elevated by Andro (**Fig. 6B, D, E**). That is to say, Andro alone was able to induce the activation of Nrf2-HO-1 signaling pathway. Indeed, Andro time-dependently increased the levels of Nrf2, with rapid Nrf2 induction within 0.5 h of Andro exposure (**Fig. 6G, H**). Importantly, the levels of Nrf2 and HO-1 in TSI-treated cells inversely correlated with that of p-MLKL (**Fig. 6B, F**), suggesting that Andro inhibited necroptosis by activating Nrf2-HO-1 signaling pathway.

**Fig. 6.**
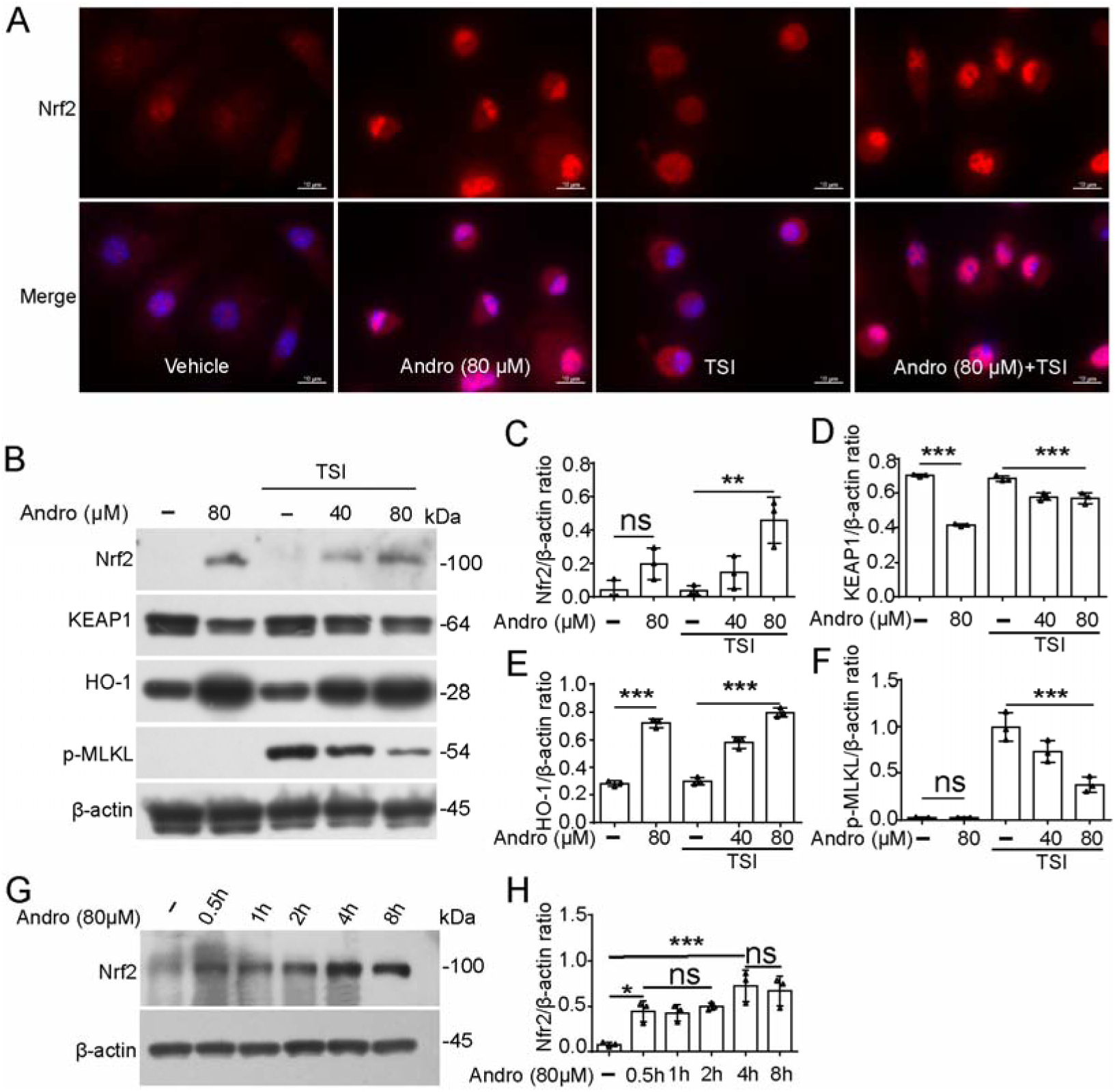
Andrographolide activates the Nrf2 signaling pathway by promoting the translocation of Nrf2 into the nucleus. BMDMs were treated as described in Fig. 1. (**A**) The distribution of Nrf2 (red) was observed by immunofluorescence microscopy. Nuclei were stained with Hoechst 33342 (blue). Scale bars, 10 μm. (**B**) Indicated proteins were analyzed by Western blotting. β-actin was used as a loading control. (**C-F**) Relative levels of indicated proteins to that of β-actin in cell lysates were determined, with the levels of the TSI group being set as 1.0, respectively. (**G, H**) BMDMs were treated with Andro for different time lengths. The levels of Nrf2 in the cell lysates were analyzed by Western blotting. Data are presented as mean ± SD (n = 3). *\*P* < 0.05; *\*\*P* < 0.01; *\*\*\*P* < 0.001; ns, not significant.

### 3.7 The derivatives of Andro without Nrf2-activation capacities do not have anti-necroptotic activity upon the necroptotic inducers

To establish structure-activity relationships, we compared Andro with its derivatives, including dehydroandrographolide (**Dehydro**), neoandrographolide (**Neo**), 14-deoxy-11,12-didehydroandrographolide (**Deoxy-didehydro**), and 14-deoxyandrographolide (**Deoxy**) (**Fig. 7A**). Andro is a labdane diterpenoid with three hydroxyl groups (-OH) at positions C-3, −14, and −19, and a double bond at C-12(13). Compared to Andro, Deoxy and Dehydro have lost the C-12(13) double bond, while Deoxy-didehydro shifts it to the position C-11(12); Neo links a glucoside moiety to its C-19 hydroxyl group (which improves water solubility) but loses the C-3 hydroxyl group; and all the four derivatives have lost the C-14 hydroxyl group. Among these chemicals, only parent Andro could profoundly activate the Nrf2-HO-1 signaling (**Fig. 7B**, **C**). Andro-treated cells showed substantially stronger nuclear Nrf2 immunofluorescence as compared to 14-deoxyandro-treated ones (**Fig. 7D**). Functional assays showed that none of the derivatives inhibited the cell death induced by LI; conversely, Deoxy, Deoxy-didehydro, and Dehydro even dose-dependently exacerbated it (**Fig. 7E-I**). These results suggest that Nrf2-HO-1 activation is essential for Andro’s anti-necroptotic activity.

**Fig. 7.**
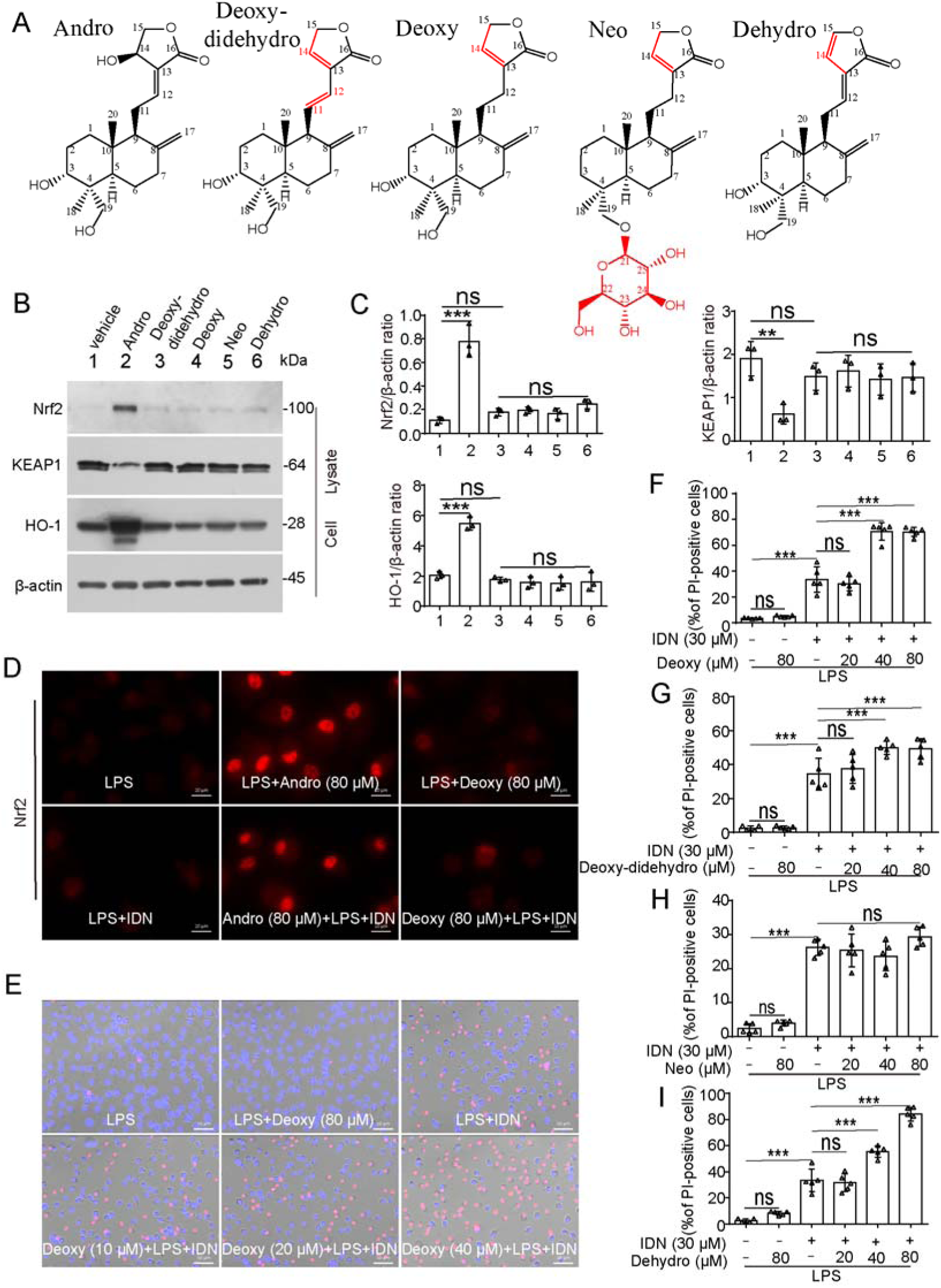
The anti-necroptotic activity of andrographolide instead of its derivatives is related to its activation of Nrf2-HO-1 signaling pathway. **(A)** Chemical structure formulas of Andrographolide (Andro), dehydroandrographolide (Dehydro), neoandrographolide (Neo), 14-deoxy-11,12-didehydroandrographolide (Deoxy-didehydr**o**), and 14-deoxyandrographolide (Deoxy). **(B, C)** BMDMs were treated with Andro or its derivatives for 8 h. Protein expression levels were analyzed by Western blotting. **(D-I)** BMDMs were pretreated with or without Andro or its derivatives for 1 h, followed by stimulation with LPS plus IDN-6556 in the presence or absence of Andro or its derivatives for 2 h. **(D)** The distribution of Nrf2 (red) was observed by immunofluorescence microscopy. Nuclei were stained with Hoechst 33342 (blue). Scale bars, 10 μm. **(E)** Cell death was evaluated through staining cells with PI (red) and Hoechst 33342 (blue). Scale bars, 50 μm. (**F-I**) The proportions of PI-positive cells relative to the total number of cells stained with Hoechst 33342 were calculated. Data are shown as mean ± SD (n = 5). *\*\*P* < 0.01; *\*\*\*P* < 0.001; ns, not significant.

### 3.8 Andro alleviates the uterus injury induced by Tumor necrosis factor-α (TNF-α) in mice

Given the established role of TNF-α in inducing necroptosis-driven inflammation ^[18]^, we evaluated the anti-inflammatory effect of Andro in TNF-α-induced systemic inflammatory response syndrome (SIRS) in mice. Histopathological analysis revealed extensive uterine necrosis in TNF-α-treated-group (**Fig. 8A**). With Andro treatment, however, the uterus injury induced by TNF-α was markedly alleviated (**Fig. 8A**). Western blot analysis showed that Andro reduced the levels of p-MLKL (**Fig. 8B, C**), confirming necroptosis suppression. These results show that Andro at low doses (< 1 mg/kg, bw) could alleviate inflammation that is associated with necroptosis.

**Fig. 8.**
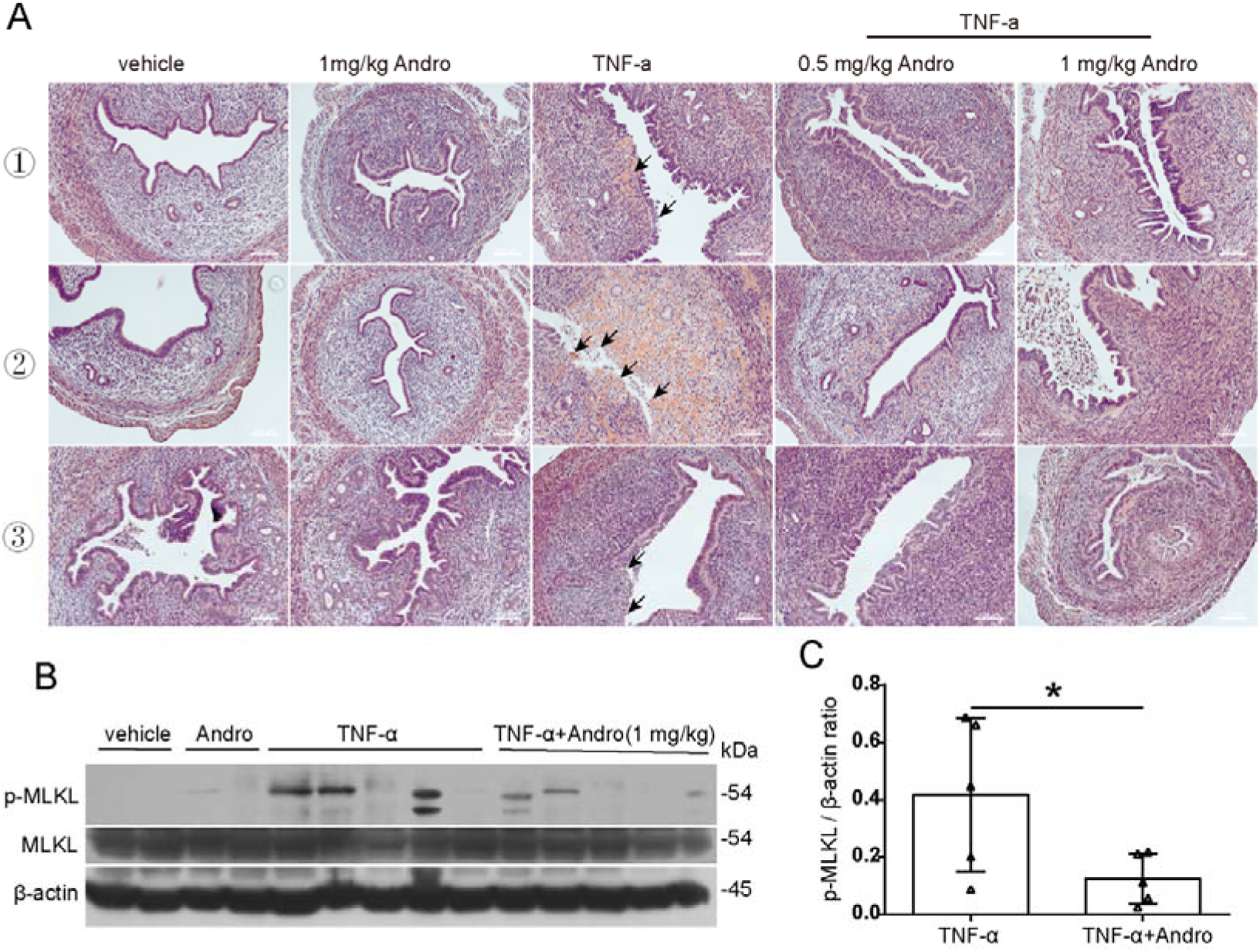
Andro mitigates the severity of TNF-α-induced uterus injury in mice. Wild-type female mice were intraperitoneally injected with sterile PBS or Andro thrice, at 1 h before, and 2 h and 5 h after the injection of TNF-α (5[μg per mouse) via the tail vein. Uteruses were collected 9[h after the challenge with TNF-α. (**A**) Representative H&E-stained histology of uteruses from the mice. Microscopic images of histological sections are shown at 100× magnification. Scale bar, 100 µm as indicated. Epithelial injury was indicated by black arrows. (**B**) Western blot analysis of p-MLKL expression in the uterus tissues from each group. The loading control was β-actin. (C) Relative levels of p-MLKL to that of β-actin was determined. An unpaired Student’s t-test was utilized to evaluate the differences between the two groups. Data are shown as mean ± SD (n = 5). \**P* < 0.05.

## 4. Discussion

Previous studies have established Andro’s dual suppressive effects on apoptosis ^[32]^ and pyroptosis ^[10, 33]^, but its potential modulatory role in necroptosis remains unknown. The findings of the current study demonstrate Andro’s broad-spectrum inhibitory capacity against LI- or TSI-induced necroptosis in macrophages and other cell types. Further investigation on the underlying mechanisms revealed that Andro blocks the RIPK1/RIPK3/MLKL pathway while upregulates Nrf2 and its downstream antioxidant targets, thus decreasing intracellular ROS and preserving MMP. These results suggest that Andro can be used as an effective inhibitor of necroptosis.

Necroptosis contributes to diverse pathological conditions, such as kidney injury ^[34]^, heart diseases ^[35]^, cancers ^[36]^, and neurodegenerative diseases ^[37]^ *etc*. Several inhibitors have been developed to target the RIPK1-RIPK3-MLKL necroptotic pathway, including Nec-1 (targeting RIPK1) and GSK’872 (targeting RIPK3) ^[38]^.

Notably, ROS-mediated cysteine oxidation in RIPK1 initiates its autophosphorylation; subsequently, p-RIPK1 recruits RIPK3 to form a necrosome, ultimately leading to necroptosis ^[25, 39]^. However, the precise mechanism, especially how ROS regulates the RIPK1-RIPK3-MLKL pathway, remains unclear. Our group’s prior studies have shown that mtROS scavenging via inhibiting reverse electron transport prevents necroptosis and PANoptosis ^[18, 40]^. Recently, our lab found that some phytochemicals, including baicalin ^[24]^ and celastrol ^[22]^, can inhibit necroptosis by suppressing mtROS. Consistent with this paradigm, Andro effectively attenuated both total ROS and mtROS while preserving mitochondrial integrity.

Previously, Andro has been reported to target Drp1, a protein that is fundamental for mitochondrial and peroxisomal fission ^[41]^. Given that RIPK1/ROS-mediated translocation of MLKL, Bak, and Drp1 to the mitochondria drives MMP collapse and finally results in necroptosis ^[29]^, Andro’s inhibition of Drp1, Bcl-2, and Bak oligomerization suggests possible cross-talk between mitochondrial fission machinery and necrosome formation, but whether it inhibits necroptosis by directly targeting Drp1 needs more investigation.

Another target of Andro that has been reported is Keap1 ^[42]^, which is a key regulator of cellular Redox homeostasis as described above ^[43]^. The Keap1/Nrf2 regulatory network extends to the thioredoxin (Trx) and glutathione (GSH) systems, which coordinately maintain the native thiol status of proteins ^[44]^. In this study, we show that the levels of Nrf2 and HO-1 proteins were increased upon Andro treatment both in the presence or absence of necroptotic inducers (TSI). Interestingly, the levels of Nrf2 and HO-1 were inversely correlated with the levels of p-MLKL, a marker of necroptosis. The concurrent inability of derivatives to activate Nrf2-HO-1 signaling or inhibit necroptosis further supports this pathway’s essential role.

While our study establishes Nrf2-HO-1 activation as a key anti-necroptotic mechanism, it is unclear how Nrf2 and HO-1 precisely regulate the formation of necrosomes. Since upregulation of Nrf2 and HO-1 by Andro has been observed in various studies^[45–47]^, we hypothesize that Andro likely targets KEAP1 to prevent ROS outbreak. Indeed, the current study showed that Andro prevented the oligomerization of some mitochondrial proteins that are critical for cell survival, such as Bcl-2 and Bak, through their disulfide bonds. These cross-linked proteins not only induced mitochondrial damage but also participated in necrosome assembly, suggesting a feedforward mechanism whereby oxidized Bcl-2/Bak both induce mitochondrial permeabilization and serve as necrosome scaffolds. Validating this hypothesis requires advanced techniques such as *in situ* crosslinking proteomics and real-time ROS biosensor imaging.

Our study also showed that Andro at low doses (≤ 1mg/kg, bw) could alleviate the uterus damage induced by TNF-α in mice. Aligning with our previous SIRS findings^[18]^, TNF-α induced necroptosis in the uterus tissues, which was significantly suppressed by Andro treatment. It is noted that Andro consistently upregulated Nrf2 and HO-1 levels irrespective of TSI co-stimulation. Although our study showed that a short-term and low-dose of Andro treatment show no cytotoxicity towards normal cells, caution is warranted since chronic/high-dose toxicity had been highlighted in comprehensive Andro pharmacological reviews^[48]^. Nonetheless, given its remarkable anti-necroptotic and anti-inflammatory properties, Andro can potentially be developed as an external medicine ^[48]^.

In this study, we showed that Andro could inhibit necroptosis, but its drivatives failed, indicating critical structure-activity relationships. All the 4 derivatives and Andro have been reported with anti-inflammatory and anti-cancer properties^[49–51]^, as well as with protective effects on hepatotoxicity induced by toxins^[52–55]^. They also have viricidal activities^[56]^, but the spectrum of viruses they can kill vary greatly. For example, Andro and Deoxy can kill Foot-and-Mouth Disease Virus 3C^pro^, but Neo cannot^[57]^. Similarly, Andro and Deoxy-didehydro significantly inhibited thrombin-induced platelet aggregation, but Neo exhibited little or no activity^[58]^. The derivatives and Andro differ in their targeting and bioavailability due to their slight structural differences such as double bonds and hydroxyl substitutions (**Fig. 7A**). Systematic comparison of derivative pharmacodynamics could illuminate optimal molecular scaffolds for pathway-specific drug design.

In summary, the results of this study suggest that Andro can inhibit necroptosis by reducing ROS, protecting mitochondria from MMP loss, and inhibiting the formation of necrosomes. However, the inability of derivatives to activate Nrf2/HO-1 signaling or inhibit necroptosis highlights Andro’s unique structural requirements. Our data imply that Andro is a potential phytochemical for treating necroptosis-related diseases.

## Conflict of Interest

All authors declare that they have no competing interest with respect to the contents of this article.

## Acknowledgement

This work was supported by Medical Joint Fund of Jinan University (No. YXJC2024001), Funding of Science and Technology Projects in Guangzhou (2024A03J0809), and the National Natural Science Foundation of China (Nos. 82274167, 81873064 and 81773965). We thank Prof. Yong-tang Zheng in Kunming Institute of Zoology, CAS, for his help in this study.

## Authors contributions

**Conceptualization:** Ouyang D. **Data curation:** Li Q, Lu N. **Funding acquisition:** Zha Q, He X, Ouyang D. **Investigation:** Lu N, Li Q, Duan L, Xu R, Shi F, Zhou Z, Gan Y, and Huang X. **Methodology:** Li Q, Li Y. **Project administration:** Ouyang D, Hu B, Li J. **Supervision:** Ouyang D, He X. **Validation:** Gan Y. **Writing**―**original draft:** Lu N, Li Q. **Writing**―**review & editing:** Ouyang D, He X.

---

**Fig. S1.**
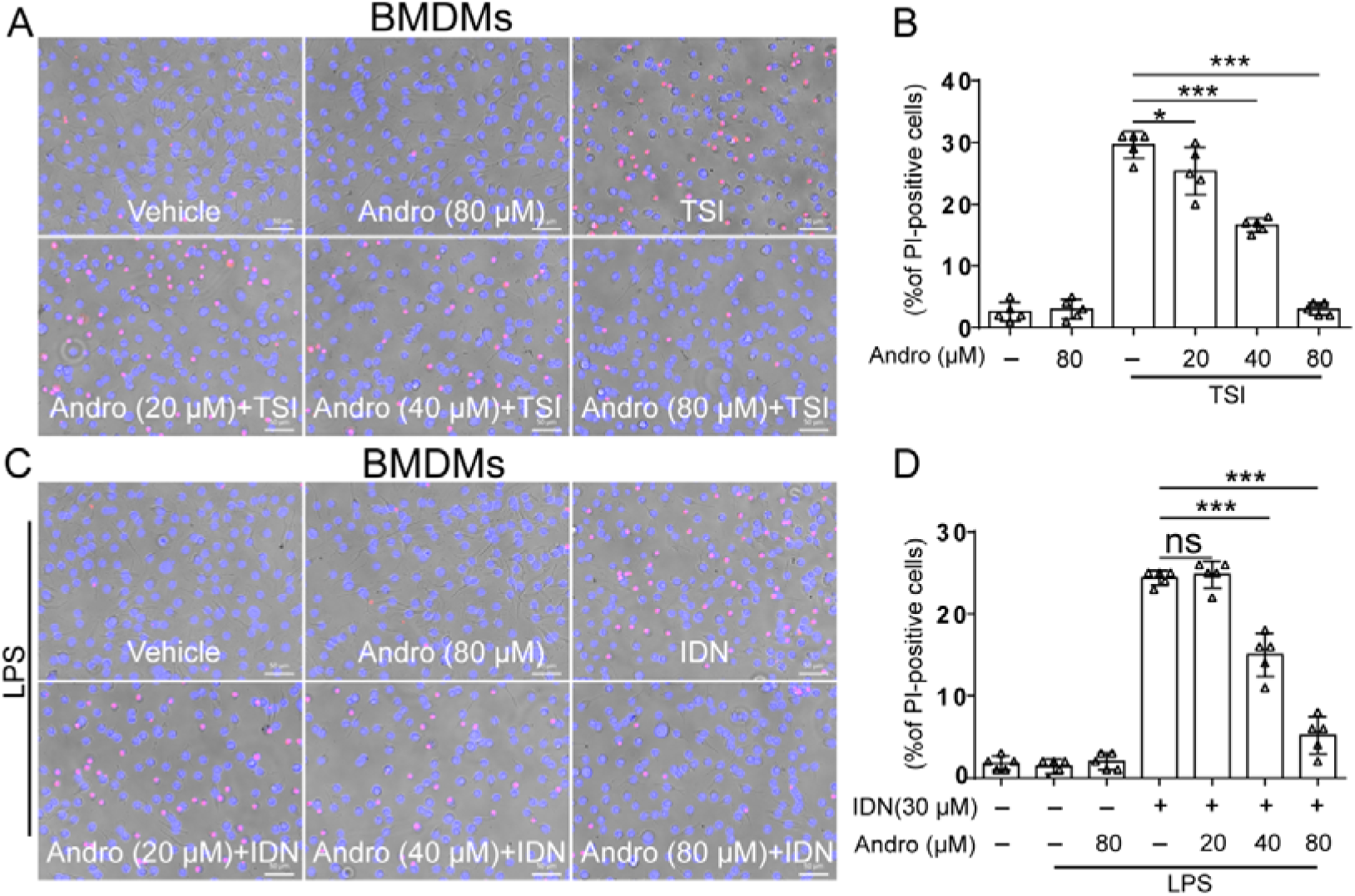
Andro suppresses necroptosis in macrophages. Bone marrow-derived macrophages (BMDMs) were pretreated with or without Andro for 1 h, followed by stimulation with the combination of TNF-α, LCL-161, and IDN-6556 (TSI) (**A, B**), or LPS plus IDN-6556 (LI) (**C, D**) in the presence or absence of Andro for 2 h. Cell death was evaluated by staining cells with PI (red, staining lytic cells) and Hoechst 33342 (blue, staining all nuclei). Scale bars, 50 μm. Data are shown as mean ± SD (n = 5). *\*P* < 0.05; *\*\*\*P* < 0.001; ns, not significant.

**Fig. S2.**
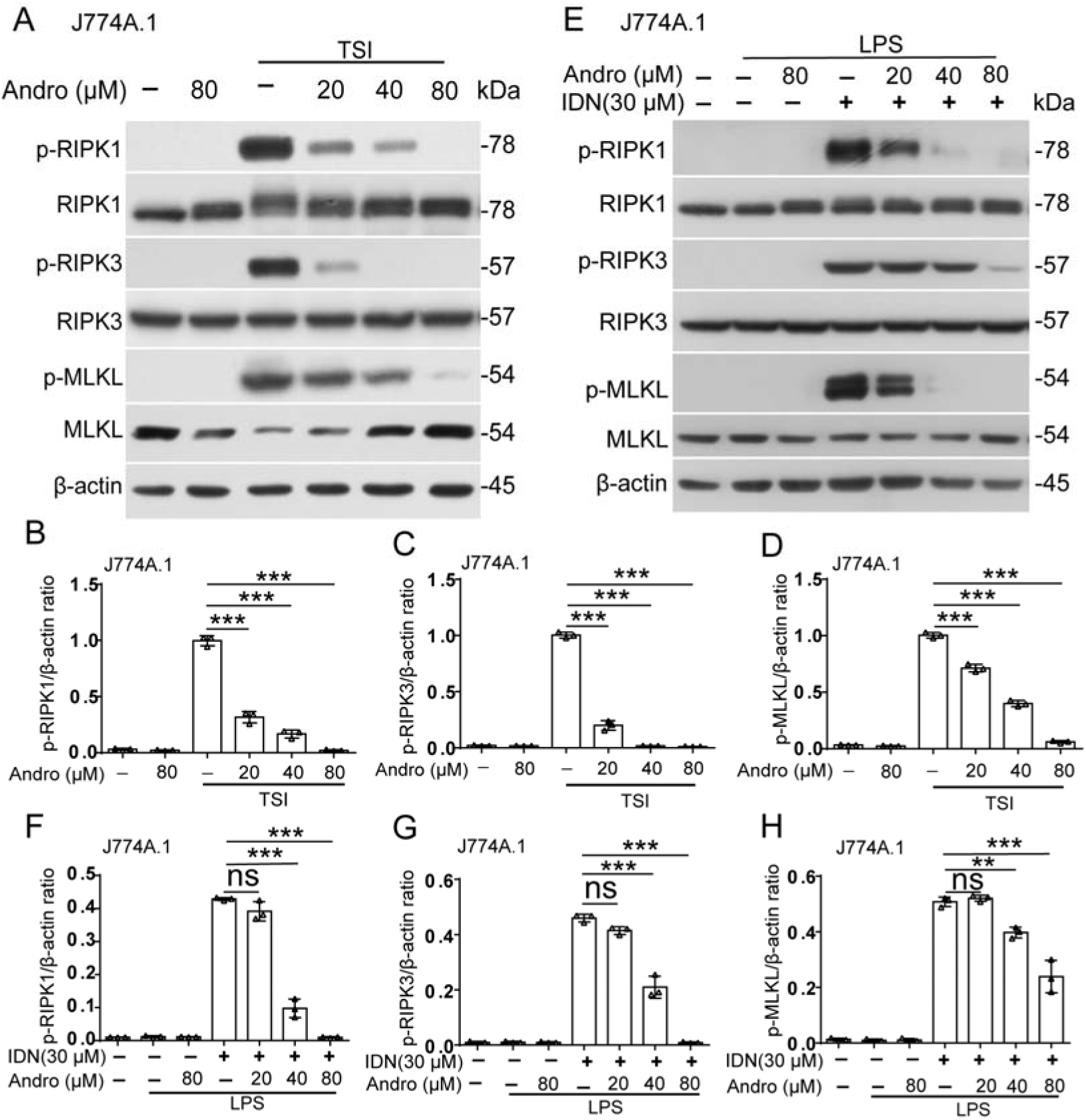
Andrographolide suppresses the necroptotic signaling pathway. J774A.1 cells were treated with TNF-α, LCL-161, and IDN-6556 (TSI) (**A-D**), or LPS plus IDN-6556 (LI) (**E-H**) as described in Fig. 1. Western blotting was used to detect the proteins of the necroptotic pathway. The loading control was β-actin. Relative levels of indicated proteins to that of β-actin in cell lysates were determined, with the levels of the TSI or LI group being set as 1.0, respectively. Data are shown as mean ± SD (n = 3). *\*\*P* < 0.01; *\*\*\*P* < 0.001; ns, not significant.

**Fig. S3.**
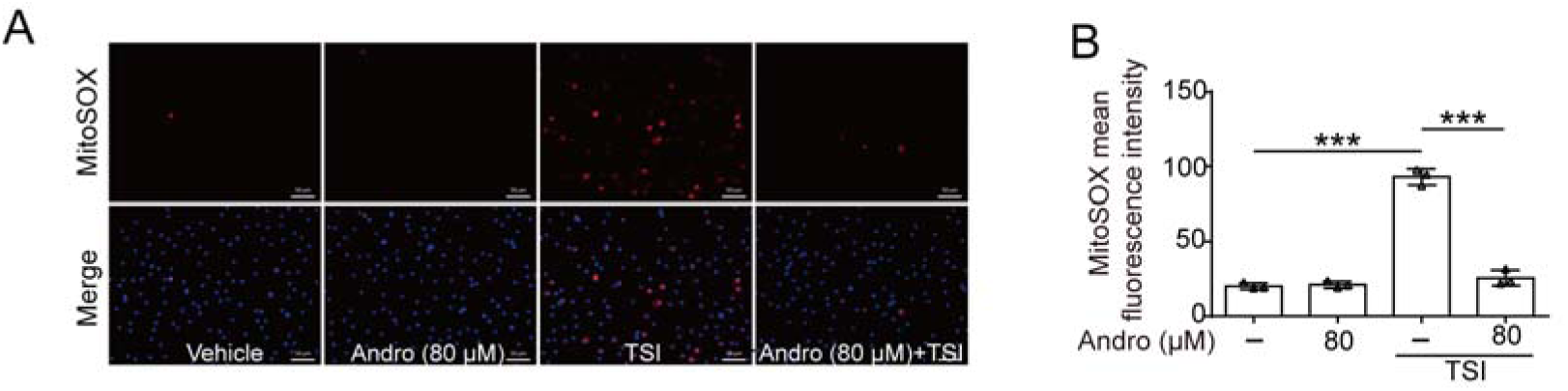
Andrographolide reduces mitochondrial reactive oxygen species (ROS) in TSI-treated macrophages. **(A)** Bone marrow-derived macrophages (BMDMs) were treated as described in Fig. 1. After staining the cells with mitoSOX, mitochondrial superoxides (red) were observed by fluorescence microscopy. Nuclei were stained with Hoechst 33342 (blue). (**B**) Ratios of cells with mitochondrial superoxides to all cells. *\*\*\*P* < 0.001.

## Notes

### Competing Interest Statement

The authors have declared no competing interest.

